# TGF-β3 Promotes Trophoblast Development via ACSS2-Dependent Permissive Lipid Metabolism

**DOI:** 10.1101/2025.01.02.631122

**Authors:** Francesca Boffa, Margherita Moncada, Martina Lo Sterzo, Luca Palazzese, Aaurora Scuderi, Marika Domenicone, Emanuele Capra, Barbari Lazzari, Luisa Gioia, Ramiro Alberio, Domenico Iuso, Pasqualino Loi, Marta Czernik

**Affiliations:** University of Teramo, Department of Veterinary Medicine, 64100 Teramo, Italy; CNR-IBBA, Institute of Agricultural Biology and Biotechnology, National Research Council, Via Einstein, Lodi 26900, Italy; School of Biosciences, University of Nottingham, Sutton Bonington Campus, Nottingham, LE12 5RD, UK

**Keywords:** Blastocyst, *In vitro*, Ovine, TGF-β3, Trophoblast

## Abstract

Transforming growth factor-beta (TGF-β) supports the in vitro maintenance of embryonic and trophoblast stem cells. Here, we demonstrated that, in a sheep embryo model, the transition from morula to blastocyst is positively regulated by TGF-β3, primarily through its promotion of trophoblast development. Our results indicate that morulae treated with TGF-β3 develop at a higher rate into blastocysts, characterized by an expanded trophoblast layer marked by CDX-2 expression. In blastocysts, TGF-β3 mediates transcriptional activation of genes involved in cell adhesion and lipid metabolism pathways, leading to remarkable in vitro outgrowth expansion and a substantial increase in trophoblast lipid droplet content. Functional analysis reveal that the positive effects of TGF-β3 are mitigated by inhibition of Acetyl-CoA Synthetase Short-Chain Family Member 2 (ACSS2), a key enzyme upregulated by TGF-β3 and a promoter of de novo lipgenensis. These findings suggest that TGF-β3 modulates lipid metabolism during blastocyst formation and may play a potential role in regulating implantation and placental development.

## Introduction

Assisted Reproduction Techniques (ARTs) are widely used in human infertility clinics, the animal industry, and fundamental research. In vitro-produced farm animal embryos serve a pivotal role not only as models for studying human fertility and embryonic development but also in enhancing reproductive efficiency, improving genetic stock, and contributing to the conservation of endangered species [Calhaz-Jorge et al., 2020].

Despite notable advances, current *in vitro* culture systems still fail to achieve similar efficiency as in vivo embryo production [Viana, 2019]. The proportion of embryos that successfully reach the blastocyst stage remains relatively low, often not exceeding 20–40% in species such as sheep [Zhu et al., 2018], and pigs [Chen et al., 2022]. A critical bottleneck in *in vitro* embryo development [Gualtieri et al., 2024] has been identified during the morula to blastocyst transition, when the inner cell mass and the trophectoderm are established. Despite the significance of this developmental step, its molecular mechanisms remain poorly unsterstood. The transcriptome profile of blastocysts revealing distinct gene expression patterns depending whether they are produced *in vivo* or *in vitro* [Gad et al., 2012]. Furthermore, it was recently reported the metabolomic and epigenetic dysfunctions lead to the arrest of *in vitro* fertilized embryos [Yang et al., 2022]. In this study, we investigated the impact of the TGF-β in the morula-to-blastocyst transition *in vitro* [Qian et al., 1992]. Previous studies reported TGF-β factors employed as key supplement in culture media for embryonic [Osnato et al., 2021] and trophoblast stem cells [Erlebacher et al., 2004].The transforming growth factor-beta (TGF-β) family is defined as a regulator of metabolism, particularly to enhance glucose uptake [Kitagawa et al., 1991, Wu & Hill, 2009] and as cytokine playing a crucial role in cell differentiation and maintenance of pluripotency [Sakaki-Yumoto et al., 2013; Gordeeva, 2019; Massagué, 2012]. TGF-β is characterized by different isoforms, encoded by distinct genes -TGF-β1, TGF-β2 and TGF-β3 – each with a molecular weight of approximately of 25kDa and sharing between 71-79% sequence identity [Baardsnes et al., 2009]. Despite their genetic differences, these isoforms exhibit a highly similar three-dimensional structure, making them nearly indistinguishable in most cell-based reporter gene and growth inhibition assays. Building on this evidence, we hypothesized the TGF-β supports blastocysts formation, potentially leading to more efficient *in vitro* embryo development resembling their *in vivo* counterparts. Our findings identify a role of TGF-β3 in enhancing in vitro blastocysts development by promoting trophoblast differentiation and adhesion through ACSS2-dependent lipid metabolism. These results suggest that TGF-β3 holds significant potential for improving outcomes in *in vitro* embryo production offering a promising avenue for enhancing embryo quality and implantation success in both animal and human applications.

## RESULTS

### TGF-β3 enhances *in vitro* blastocyst development

Our first objective was to determine the effects of TGF-β1 and TGF-β3 (0.5 ng/ml) on embryo quality after in vitro fertilization. Embryos were treated from day 4 (morula stage) to day 7 of embryo culture. Although TGF-β1 did not enhance the blastocysts development (Table 1), supplementation with TGF-β3 led to significant increase of blastocysts rates (45.8% ± 12.5 vs. 28.6% ± 11.8) compared to CTR (Chi-square; P <0.0001) (Table 1, Fig. 1A, B and C).

**Table 1.** Effect of TGF-β supplementation on embryo development. CTR (n=17), TGF-β1 (n=3), TGF-β3 (n=17). The significant difference is related to the same column (Chi-square Test; ^****^ indicate a P<0.0001).

| Group | 2cells/Oocytes (%) | Blastocysts/2cells (%) | Hatched at D7/Total blastocysts (%) |
| --- | --- | --- | --- |
| CTR | 379/685 ($55.3\% \pm 17.2$) | 100/379 ($28.6\% \pm 11.8$) | 16/100 ($16\% \pm 0.8$) |
| **TGF-β1** | 205/376 (56.7% ± 15.3) | 51/205 (25.6% ± 15.3) | 9/51 (17.6% ± 2.4) |
| **TGF-β3** | 395/685 (57.6% ± 15.2) | 181/395 (45.8% ± 12.5) | 79/181 (43.6% ± 4.6)**** |

**Figure 1.**
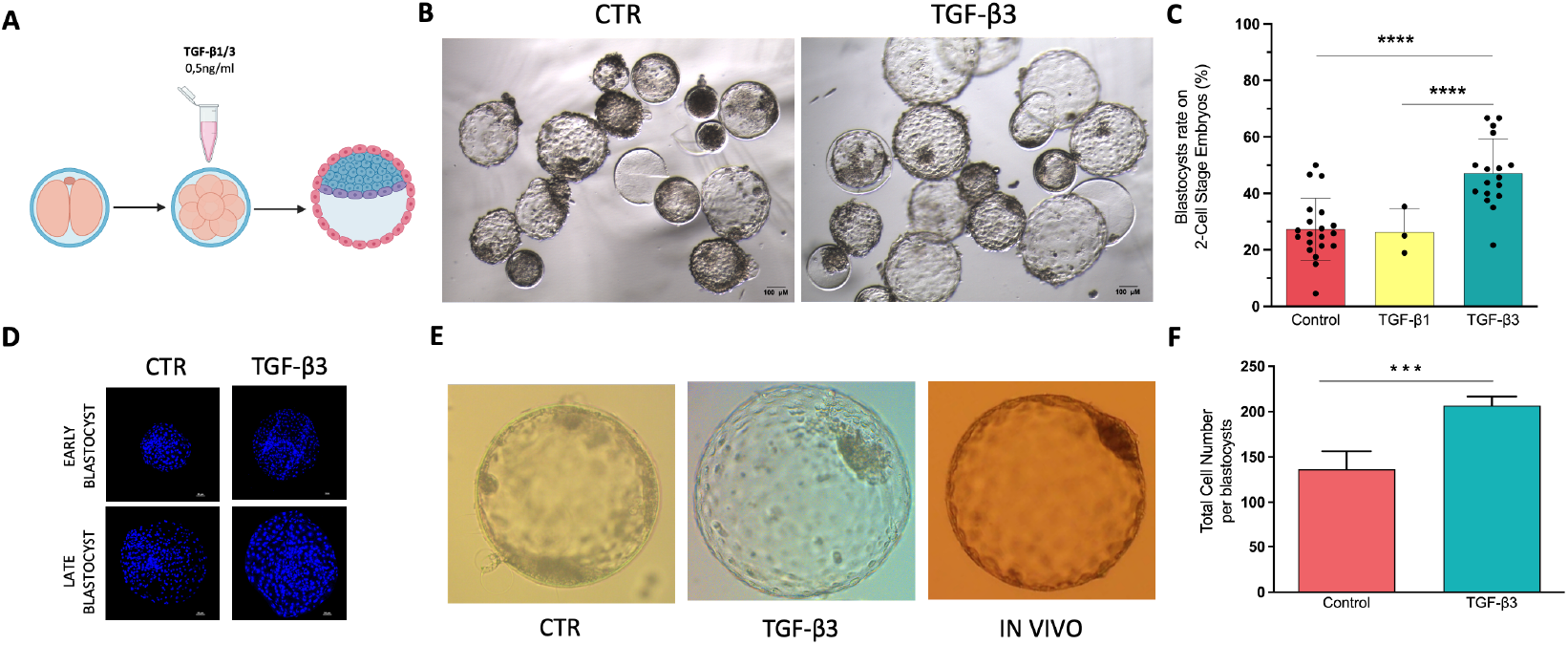
Effect of the treatment with TGF-β1 and TGF-β3 on in vitro embryo development. (A) Schematic model illustrating the experimental design; (B) Representative brightfield images of D7 embryos from the control group (CTR) and embryos cultured in presence of TGF-β3, (C) Blastocysts rate (%) on 2-cell stage embryos. ^****^ indicate significant differences (Chi-square Test; P<0.0001). Bars represent % mean± standard deviation. Every dot represents the replicates involved in the analysis, (D) Representative images of nuclear-stained D7embryos at two different developmental stages, early and late blastocysts, for control (CTR) and TGF-β3 treated groups, (E) Representative images of embryos from CTR, TGF-β3, and in vivo recovery groups at equivalent developmental stages, (F) Graph presenting the Total Cells Number of cells per blastocysts n CTR and TGF-β3 groups. ^***^ indicate significant differences (n= 10 for each group)(Unpaired t test; P<0.001) Bars represent mean ± s.d

Furthermore, blastocysts treated with TGF-β3 exhibited significally higher cell count (CTR: 136.4 ± 8.86 vs TGF-β3: 206.6 ± 4.53; means ± d.s.; P<0.001; Fig. 1D, F) and demonstrated improved morphological quality, as evidenced by a higher percentage of hatched embryos on day 7 (Table 1).

### TGF-β3 treatment promotes trophoblast expansion in vitro

Immunofluorescence analysis for CDX2 and OCT4 revealed a significant increase in the number of CDX2-positive cells, which marks the trophectoderm layer, in the TGF-β3 treated blastocysts compared to the control group (CTR: 95.3 ± 33.8 vs TGF-β3: 204.5 ± 78.3; means ± d.s.; P<0.01; Fig. 2A, B). In contrast, the number of OCT4-positive cells, marking the epiblast, remained unchanged (CTR: 13.6 ± 2 vs TGF-β3: 13.2 ± 2; means ± d.s.; Fig. 2A, B). However, despite the observed increase in trophectoderm cell numbers, transcriptomic analysis did not reveal any significant upregulation of lineage-specific gene expression associated with the three major lineages of the embryo at this stage of development (Fig. 2C). This suggests that while TGF-β3 promotes trophectoderm proliferation, it does not appear to influence the molecular differentiation pathways governing lineage specification in blastocysts. (Fig. 1E, 2B) [Li et al., 2021; Nilsen-Hamilton, 1990].

**Figure 2.**
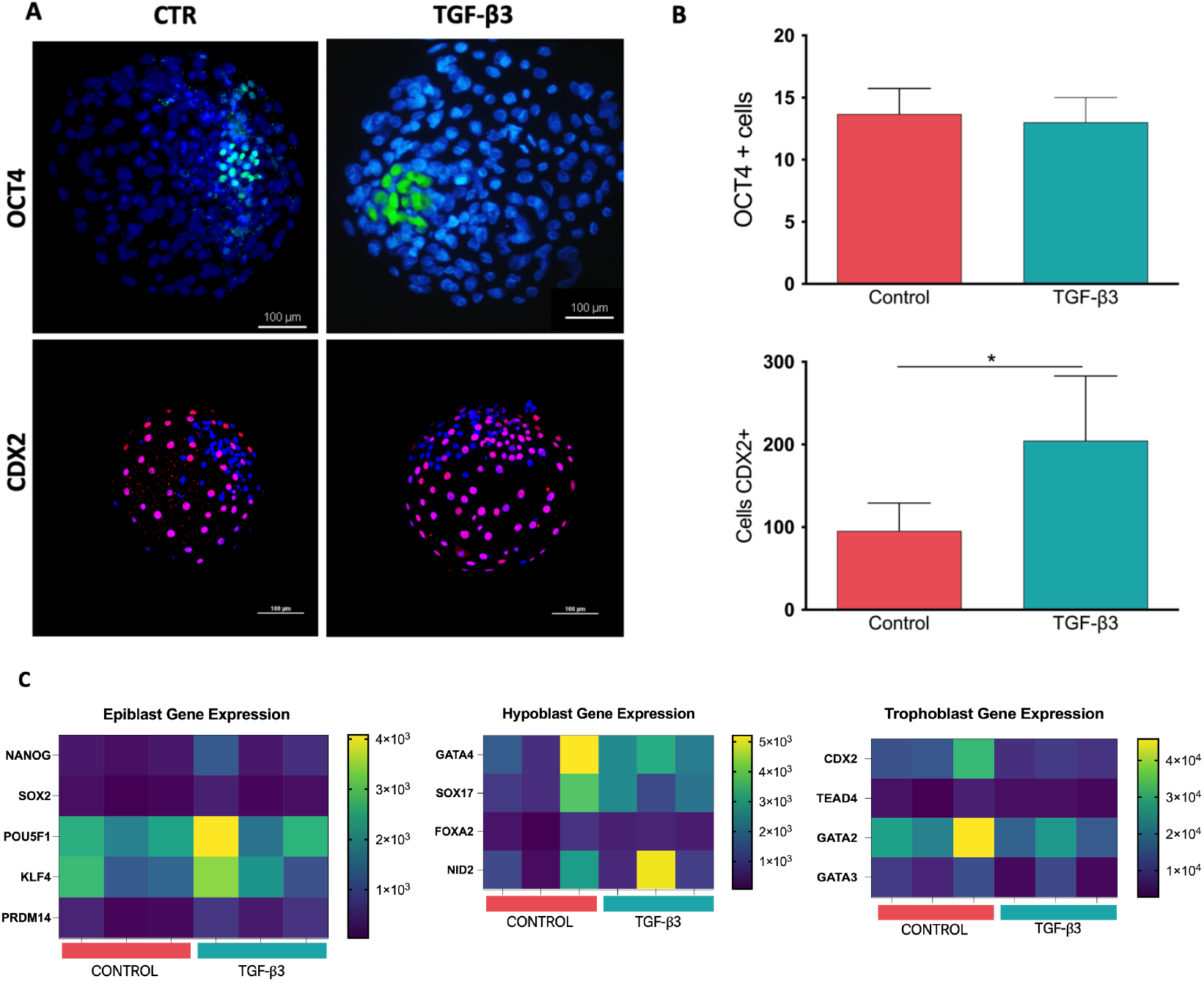
Epiblast and Trophoblast development in D7 embryos. (A) Representative immunostaining of OCT4, a hallmark gene of the Inner Cell Mass (ICM), and CDX2, a marker indicative of Trophoblast cells, in control (CTR) and TGF-β3-treated blastocysts at day 7. (B) Quantification of the number of cells positive for OCT4 and CDX2 (n=6 for each group)(Unpaired t test; p ≤ 0.01). (C) Heatmap of expression levels of selected lineage-specific genes (log2-normalized gene counts)(n=3 replicates of single blastocysts for each group).

### Transcriptome analysis of the TGF-β3 treated blastocysts

A total of 638 million paired-end reads were obtained from all samples (3 CTR, 3 TGF-β3), averaging 106.4 million ± 33.6 million reads per sample. After quality filtering, an average of 91.7% of high-quality paired reads were successfully mapped to the sheep reference genome ARS-UI_Ramb_v2.0. Generalized PCA, considering a total of 21,303 unique genes, revealed that gene expression varied after TGF-β3 treatment, with a complete separation between CTR and TGF-β3 samples along PC2, which explained approximately 23.9% of the variance. A total of 144 differentially expressed genes (DEGs) were identified in TGF-β3-treated blastocysts, with the majority being upregulated (n=129) and only fifteen downregulated compared to CTR (Fig 3A). To gain insights into the biological response of blastocysts following treatment, a functional enrichment analysis was performed on the DEGs. Notably, the expression of BMP1 [Akiyama et al., 2024], THBS1 (Thrombospondin-1) [Murphy-Ullrich & Suto, 2018], PDGF-C [Charni Chaabane et al., 2014], and LTBP1 (Latent Transforming Growth Factor Beta Binding Protein 1) [Troilo et al., 2016]—all intricately linked to or regulated by the TGF-β signaling pathway—strongly indicates that the TGF-β pathway was effectively activated in our system (Fig. 3B).

**Figure 3.**
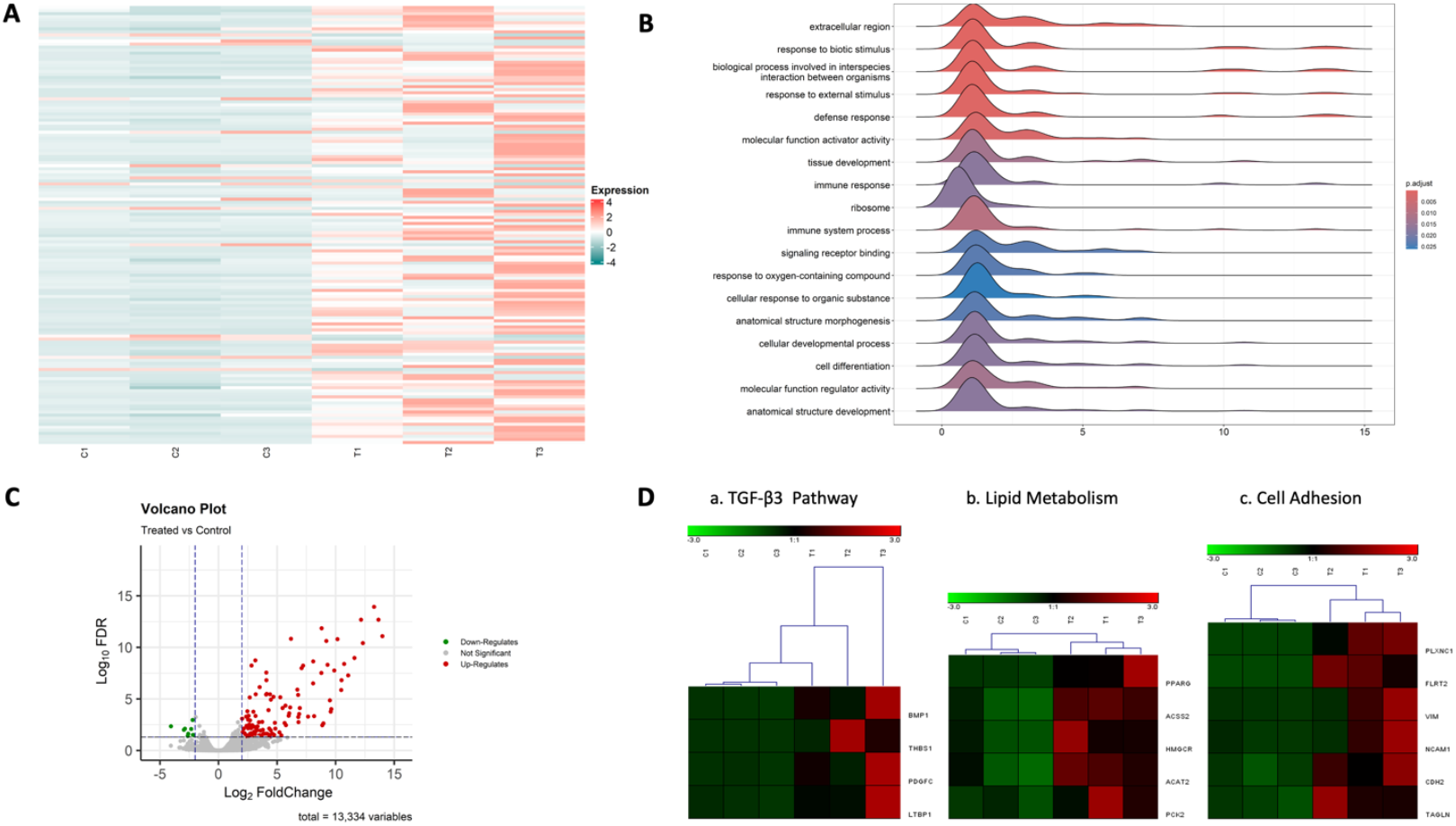
Transcriptome analysis. (A) Heatmap displaying 144 differentially expressed genes (DEGs) for each group (n=3). (B) Ridgeline plots depicting significantly enriched KEGG pathway gene sets (padj < 0.01) among DEGs in the TGF-β3-treated group. (C) Volcano plot illustrating significant DEGs with Log2 fold change and padj < 0.01. (D) Classification of key gene groups within TGF-β3-treated DEGs, associated with: (a) TGF-β signaling, (b) lipid metabolism, and (c) cell adhesion.

Moreover, when focusing on the pluripotency gene network between the groups, we observed no significant differences in the expression of KLF4, MYC, NANOG, OCT4 (POU5F1), and SOX2 between the blastocysts from both groups (Fig. 2C). This suggests that while TGF-β3 promotes the embryo formation and leads to an increase in morula development (approximately 15-20% more morulae), it maintains the same level of pluripotency and guides their development into blastocysts (Fig. 1C) [David & Massagué, 2018].

Two groups of overexpressed genes cover functional expression clusters that caught our attention. The first group formed by PLXNC1 (Plexin C1) [Ohta et al., 1995], FLRT2 (Fibronectin Leucine Rich Transmembrane Protein 2) [Seiradake et al., 2014], VIM (Vimentin) [Ivaska et al., 2007], NCAM1 (Neural Cell Adhesion Molecule 1) [Lu et al., 2002], CDH2 (Cadherin 2) [László & Lele, 2022; de Plater et al., 2024], TAGLN (Transgelin) [Wang et al., 2022] related to blastocyst and cells adhesion, represents the most upregulated genes in TGF-β3 blastocysts [Huynh et al., 2011]. The second group, including PPARG (Peroxisome Proliferator-Activated Receptor Gamma), ALDH1L2 (Aldehyde Dehydrogenase 1 Family Member L2), ACSS2 (Acyl-CoA Synthetase Short Chain Family Member 2), HMGCR (3-Hydroxy-3-Methylglutaryl-CoA Reductase), ACAT2 (Acetyl-CoA Acetyltransferase 2), e PCK2 (Phosphoenolpyruvate Carboxykinase 2, Mitochondrial) suggested that the lipid metabolism pathways are significantly increased in TGF-β3 blastocysts [Liu & Chen, 2022].

### Effect of TGF-β3 treatment on blastocyst outgrowth

*In vitro* blastocyst outgrowth is a reliable model to predict implantation potential *in vivo* [Jenkinson & Wilson, 1973]. Results showed that TGF-β3-treated blastocysts exhibited significantly higher attachment rates and outgrowth expansion compared to controls (CTR) (5/16 [31.2%] in CTR and 7/14 [50%] in TGF-β3) (Fig. 4A-B). On day 4 after the initiation of outgrowth, we measured the growth rate by calculating the area of both CTR and TGF-β3-treated embryos. We determined a significantly larger area in TGF-β3-treated blastocysts compared to controls (CTR mean: 6.4 x10^6^ µm ± 3.1 x10^6^ vs TGF-β3 mean: 8.8 × 10^6^ µm ± 0.6 x10^6^) (Fig 4B). The predominant cells observed in these cultures displayed epithelial-like morphology, characterized by dark cytoplasm and distinct cell boundaries.To confirm transcriptomic analysis indicating that TGF-β3 treatment may affect lipid metabolism, we performed Oil Red O staining. As shown in Fig. 5C and D, the TGF-β3 group contained a significantly higher number of lipid droplets compared to the control group, suggesting enhanced lipid metabolic activity (Fig. 4C, D).

**Figure 4.**
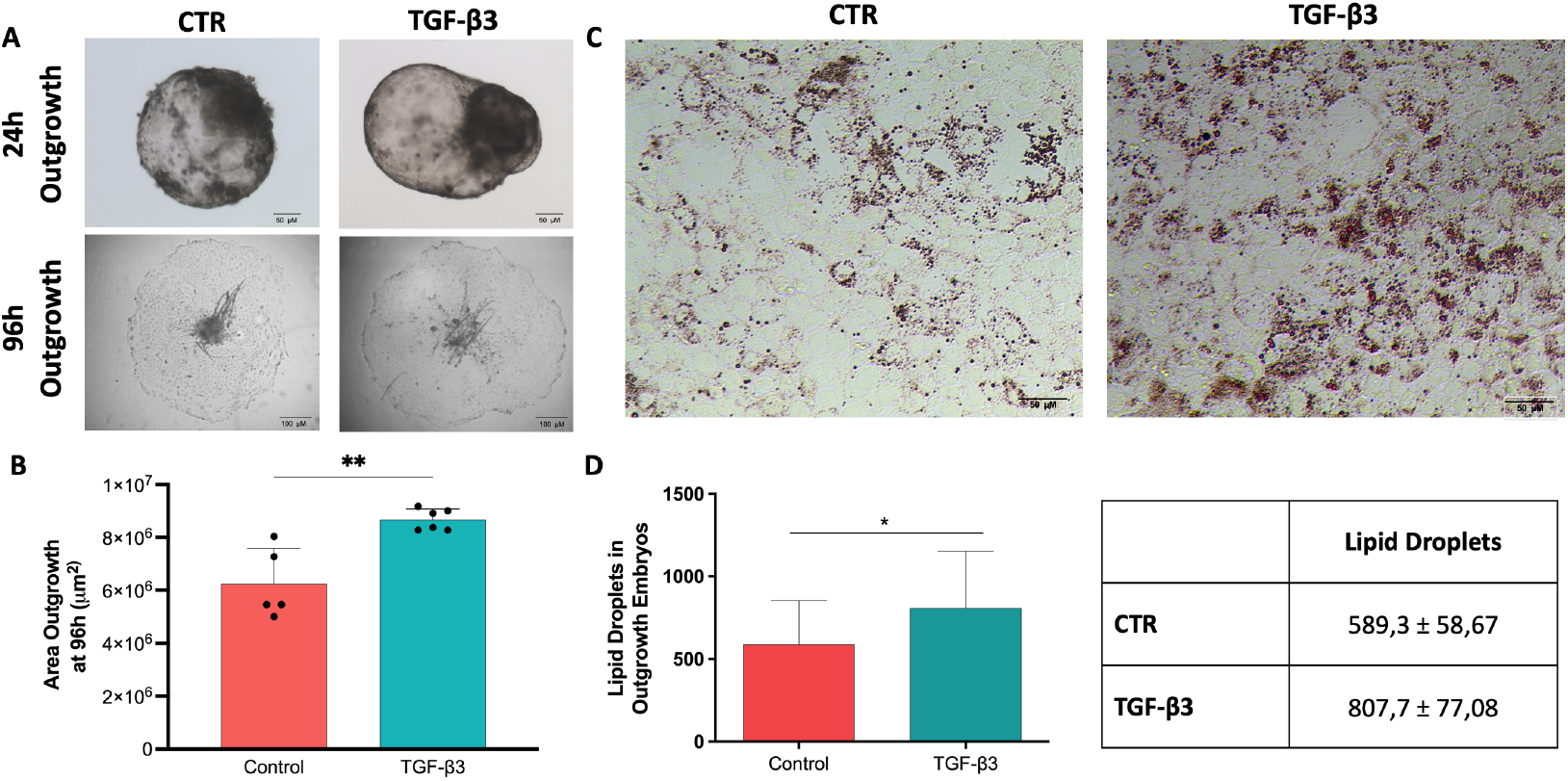
Outgrowth expansion and Lipid Metabolism involvement. (A) Representative images of blastocyst outgrowth at 24 hours post-attachment to a gelatin-coated plate and at 96 hours. (B) The outgrowth area at 96 hours was measured in μm^2^ for blastocysts cultured in outgrowth medium. Data were analyzed using Fiji software and are presented as the mean ± standard deviation (s.d.). (^**^) indicate significant differences (Kolmogorov-Smirnov test; p ≤ 0.009). Every dot represents the replicate involved in the analysis. (C) Lipid deposition in outgrowth tissues was visualized using Oil Red O staining at days 13 and 14. (D) Lipid droplets were quantified using Fiji software (ImageJ), with results shown as the mean ± s.d. (*) indicates significant differences (Unpaired t test; p ≤ 0.03).

**Figure 5.**
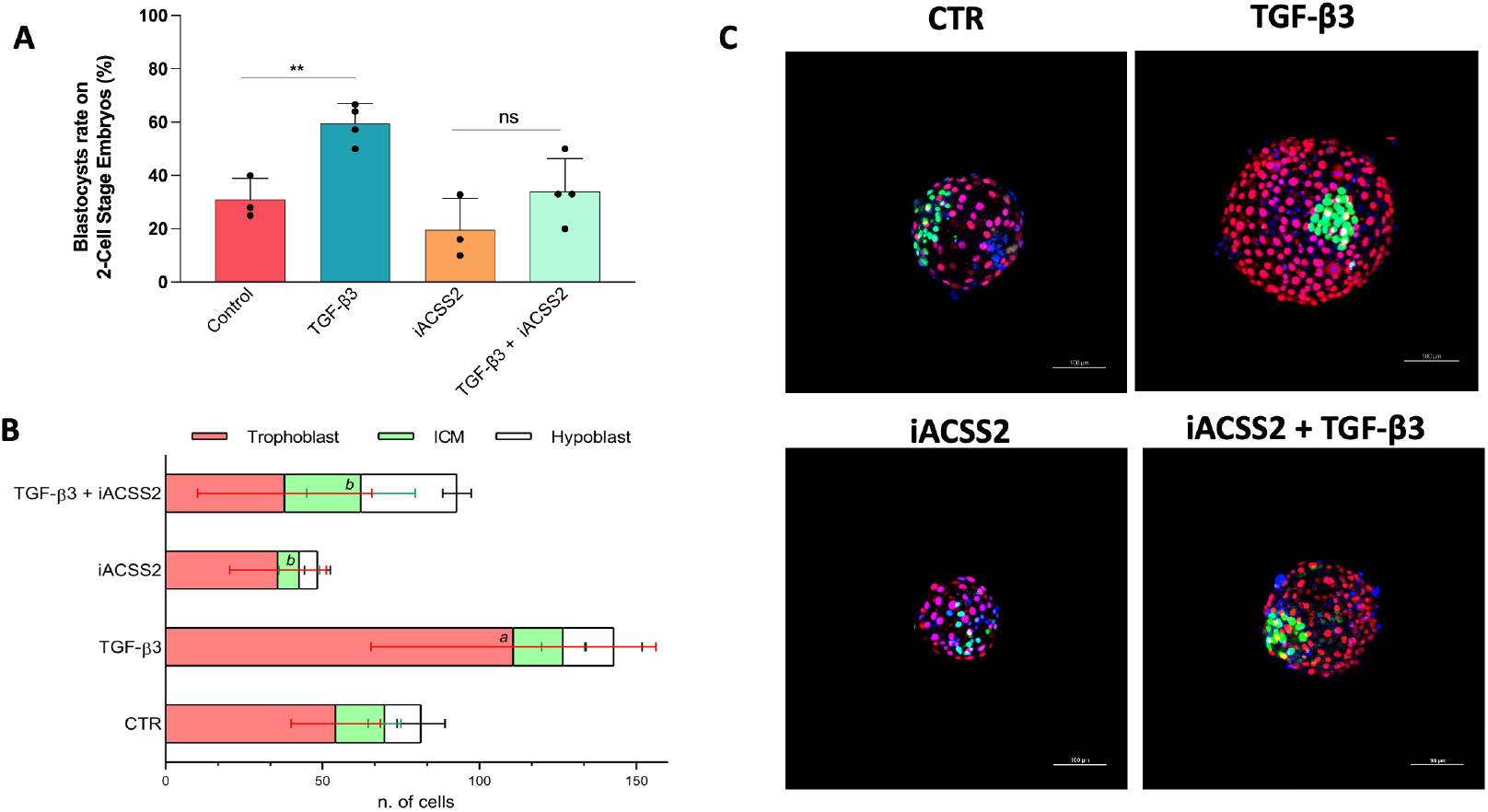
Impact of lipogenesis inhibition on TGF-β3 induced blastocysts development. (A) Blastocysts rate (%) on 2-cell stage embryos. ^*^ indicate significant differences (Chi square test; P<0.0001). Bars represent mean± s.d. Every dot represents the replicates involved in the analysis, (B) Quantification of the number of cells positive for SOX2, CDX2 and the no marked cells identified as hypoblast cells (Chi square test; P<0.0001) a= Trophoblast positive cells significantly different from all the other conditions, b= ICM positive cells significantly different in TGF-β3-iACSS2 compared to CTR-iACSS2, Bars represent mean± s.d. (C) Representative immunostaining of SOX2 (in green), CDX2 (in red), Nuclear Staining (DAPI) in the 4 groups involved with blastocysts at day 7.

### TGF-β3 promotes Blastocyst Development Through ACSS2 Activity

To determine whether the improvement observed with TGF-β3 treatment was directly related to de novo lipogenesis, as suggested by the transcriptome analysis, we inhibited key metabolic pathways at the morula stage. Specifically, we blocked ACSS2 activity, which is responsible for producing Acetyl-CoA from Acetate in lipid metabolism and its expression was upregulated in TGF-β3 treated blastocysts (Fig. 3D). The results showed that the positive effects of TGF-β3 on blastocyst development were significantly reduced by the metabolic inhibitors (CTR 31%, TGF-β3 59,4%, CTR-iACSS2 19%, TGF-β3-iACSS2 34%) (Fig. 5A). This suggests that the enhancement of blastocyst development by TGF-β3 is mediated by ACCS2 activity guiding the lipid metabolism.To further examine epiblast and trophoblast expansion in these embryos, we stained them with SOX2 (CTR 15.6 ± 5.2, TGF-β3 14 ± 6.8, CTR-iACSS2 6.8 ± 6.4, TGF-β3-iACSS2 24.3 ± 15.3), as a pluripotency and epiblast marker, and CDX2 (CTR 54.2 ± 14.2, TGF-β3 96.9 ± 45.3, CTR-iACSS2 35.8 ± 15.2, TGF-β3-iACSS2 38 ± 27.8), as a trophoblast marker [Yu et al., 2024] (Fig. 5B,C).

## DISCUSSION

In this study, we show that TGF-β3 enhances sheep blastocyst development *in vitro*. The TGF-β/activin/nodal signalling pathway is critical for pattern formation and differentiation, particularly during the pregastrulation and gastrulation stages of mammalian development [Zinski et al., 2018]. TGF-β is produced in the uterus during embryo development, coinciding with blastocyst development [Li, 2014]. While TGF-β isoforms are typically similar in most cell-based assays gene expression and growth inhibition, TGF-β2 has been noted as an exception, exhibiting 100-1000 time lower potency than TGF-β1 and TGF-β3 in cell lines lacking TβRIII [Huang et al., 2014; Cheifetz et al., 1990].

Our pilot screening using TGF-β1 and TGF-β3 showed TGF-β3 significantly improved embryo progession from the morula stage. These blastocysts closely resembled the cell number reported from in vivo-produced ones (160-180). In fact, TGF-β3 treatment increased the number of CDX2-positive TE cells in blastocyst, while the number of OCT4-positive epiblast cells remained unchanged. Blastocysts generated by TGF-β3 treatment exhibited a balanced mRNA expression of key markers for the epiblast (SOX2, OCT4, NANOG, KLF4, PRMD14), trophoblast (CDX2, TEAD4, GATA2, GATA3), and hypoblast (SOX17, GATA4, FOXA2, NID2), as confirmed by bulk RNA-seq analysis, closely mirroring control blastocysts. Furthermore, immunostaining for OCT4 and CDX2, along with gene expression analysis, revealed no significant differences also in the protein levels of these lineage markers. We conclude that TGF-β3 treatment at day 4 of *in vitro* culture supports the progression of morula to blastocysts by promoting the TE lineage.

Among all the genes highlighted in the blastocysts transcriptome analysis, Cadherin 2 and Vimentin stand out as key regulators of cell-cell adhesion. Cadherin 2 plays an essential role in establishing strong cell-cell adhesion for maintaining the integrity of the developing embryo, facilitating the morphogenetic movements that shape the blastocyst and its implantation [Ismagulov et al., 2021; Shahbazi, 2020; László & Lele, 2022; de Plater et al., 2024]. Similarly, Vimentin, an intermediate filament protein, is highly expressed in mesenchymal cells, supporting processes as cell migration, adhesion, and maintenance of cell shape, all of which are vital during embryonic development [Ivaska et al., 2007; Dave & Bayless, 2014]. The upregulation of other genes associated with blastocyst adhesion, such as Plexin C1 [Ohta et al., 1995], Fibronectin Leucine-Rich Transmembrane Protein 2 [Seiradake et al., 2014], Neural Cell Adhesion Molecule 1 [Ivaska et al., 2007], and Transgelin [Lu et al., 2002], along with enhanced TE development, supported the hypothesis that TGF-β3 -treated blastocysts may exhibit improvement in adhesion and potentionally in implantation success.

To validate this, we performed outgrowth experiments to simulate the *in vivo* attachment to uterus. The results confirmed that treated blastocysts exhibited an adhesion capacity three times greater than control blastocysts indicating a higher likelihood of successful peri-implantation development.

Furthermore, the analysis of the differentially expressed genes in TGF-β3-treated blastocysts revealed higher expression of key genes associated with lipid metabolism such as PPARG, ACSS2, PCK2 and ALDH1L2. Peroxisome Proliferator-Activated Receptor Gamma (PPARG) is reported as a key transcription factor regulating genes involved in lipid biosynthesis [Walczak & Tontonoz, 2002] and was renotably overexpressed during the elongation blastocysts phase of sheep conceptus development [Barak et al., 2008; Brooks et al., 2015, da Fonseca Junior et al., 2023].

Its increase in TGF-β3 trophoblast cells was further emphasized by the notable accumulation of lipid droplets in TGF-β3-treated trophoblast cells [Degrelle et al., 2011]. Genetic ablation of PPARG in mice results in embryonic lethality due to placental defects, which can be rescued by a wild-type placenta [Hewitt et al., 2006; Barak et al., 1999]. Moreover, reducing PPARG activity has been associated with preeclampsia [Lai et al., 2024], and consistently, in ruminant, the knockout of embryonic PPARG leads to a reduction in the total cell number in D7.5 blastocysts [McGraw et al., 2024]. Despite challenges in achieving the elongation blastocyst stage in ruminants *using in vitro* culture systems, our findings suggest that PPARG expression with the activation of lipid metabolism is a crucial pathways for blastocyst progression and elongation.

Among the overexpressed genes related to de novo lipogenesis, in combination PPARG, our focus was on ACSS2, which has been reported to be hyperactivated by TGF-β signaling (Zhu et al., 2023). ACSS2 has previously been described as a principal producer of Acetyl-CoA from acetate during embryo development and in trophoblast stem cells under metabolic stress (Zhou et al., 2021; Yu et al., 2024). The inhibition of ACSS2 decreased blastocyst development and mitigated the positive effects of TGF-β3 reducing its ability to enhance trophoblast lineage expansion, as shown by a marked depletion in CDX2-positive cell numbers. When *de novo* lipogenesis support declined, the beneficial outcomes achieved with TGF-β3 treatment were substantially reduced, failing to replicate the developmental progression observed in the untreated controls. Despite progress, the underlying mechanisms regulating these metabolic pathways remain to be fully elucidated.

Our study underscores the pivotal role of TGF-β3 in supporting in vitro blastocyst development by facilitating trophoblast development driven by ACSS2-dependent lipid metabolism. While these findings suggest TGF-β3 is a valuable candidate for advancing in vitro embryo production, further in vivo studies are required to evaluate its direct impact on peri- and post-implantion development.

## MATERIAL and METHODS

### Animals and ethics approval

All animal experiments (semen collection) have been approved by the Italian Ministry of Health, upon the presentation of the research description prepared by the ethics committee of the Istituto Zooprofilattico Sperimentale di Teramo (Prot. 944F0.1 del 04/11/2016). The number of the authorization granted by the Italian Ministry of Health is no 200/2017-PR. All methods were performed in accordance with the relevant guidelines and regulations of the Italian Minister of Health.

### Embryo production

#### Oocyte Collection and In Vitro Maturation (IVM)

Sheep ovaries were ethically sourced from local slaughterhouses and immediately transported to the laboratory within a 1-2 hours timeframe, maintaining a temperature of 37°C. Oocytes were aseptically extracted using 21 G needles in the presence of H-199 medium, which was supplemented with Bovine Serum Albumin (BSA), Gentamicin solution, and heparin. Oocytes possessing a minimum of 2-3 layers of compact cumulus cells were meticulously selected for in vitro maturation (IVM). Maturation was conducted in bicarbonate-buffered TCM-199 medium (Gibco) containing 2 mM glutamine, 0.3 mM sodium pyruvate, 100 μM cysteamine, 10% fetal bovine serum (FBS) (Gibco), 5 μg/ml follicle-stimulating hormone (FSH), 5 μg/ml luteinizing hormone (LH), and 1 μg/ml estradiol [Anzalone et al., 2016]. The IVM process occurred in 4-well culture plates, with each well containing 0.5 ml of IVM medium, incubated in a humidified atmosphere at 38.5°C with 5% CO2 for 22 hours. Following IVM, only carefully selected MII oocytes exhibiting expanded cumulus and normal morphology were employed for In Vitro Fertilization (IVF) [Anzalone et al., 2021].

#### In Vitro Fertilization (IVF) and IVC

Twenty-two hours after IVM, cumulus-oocyte complexes (COCs) displaying cumulus cell expansion were exclusively chosen for in vitro fertilization (IVF). These COCs were gently transferred into a solution containing 300 U/ml of hyaluronidase dissolved in TCM-199, washed twice in H199, and then placed in 50 μl droplets (with 8 to 10 oocytes per drop) of IVF medium (SOF-supplemented with 20% oestrus sheep serum and 16 mM isoproterenol), which were covered with mineral oil. A single straw of frozen semen, containing 100 × 10^6^ spermatozoa, was rapidly thawed in 35-37°C water and subjected to centrifugation in sperm-wash medium (bicarbonate-buffered synthetic oviductal fluid (SOF-) with 0.4% (w:v) fatty-acid-free BSA) at 1200 rpm for 5 minutes. The supernatant was discarded, and 5 × 10^6^ spermatozoa were added to each drop, followed by incubation in a humidified environment at 38.5°C, 5% CO_2_, and 7% O_2_ [Anzalone et al., 2021]. Within the initial 4 hours following IVF, meticulous oocyte cleansing was carried out by gentle pipetting in a solution of synthetic oviductal fluid (SOF-) enriched with 2% (v:v) basal medium Eagle essential amino acids (EAA), 1% (v:v) minimum essential medium (MEM)-nonessential amino acids (NEAA) (Gibco), 1 mM glutamine, and 8 mg/ml fatty acid-free BSA (SOF-aa). This process aimed to remove granulosa cells. Subsequently, the oocytes underwent three additional rinses in an in vitro culture (IVC) medium before being grouped in culture drops, with each group containing five oocytes. On day 7/8 of culture, the assessment of blastocysts was carried out using a Nikon Eclipse Ti2-U inverted microscope, aided by the Octax EyeWare Imaging Software (version 2.3.0.372).

#### Experimental Design

On day 4 of culture defined as day 0 from fertilization, embryos at the compact morula stage were divided into two experimental groups. One group was treated with TGF-β3 or TGF-β1 at a concentration of 0,5 ng/mL, which was integrated into the in vitro culture (IVC) medium. This concentration was determined based on the preliminary, pilot experiments and CL50, ensuring that the experimental conditions were optimized for obtaining and reliable results [Lyra-Leite et al., 2023]. The second group of embryos were culture in a standard IVC culture medium without TGF-β3 and serve as the control (CTR) for all subsequent experiments.

#### Embryo evaluation and total cell count

Embryos were eassessed on day 7 of culture, selecting those that had successfully hatched by this point to ensure the selection of the most advanced and viable embryos. After fixation, the total cell count of each embryo was determined using DAPI staining to visualize the nuclei. Quantification of the total number of cells was performed using QuPath software version 0.5.1, which allowed for precise cells counting.

#### Immunofluorescence

Embryos were fixed in 4% paraformaldehyde for 15 min, washed in PBS with 1% bovine serum albumin (BSA), permeabilized in 1% Triton X-100 in PBS for 15 min at room temperature (RT) and blocked in 5% BSA and 1% Tween20 in PBS for 1h at RT. Then, they were incubated overnight at 4°C with primary antibodies (S.T.1). After four washes in 1% BSA in PBS, embryos were incubated in the appropriate secondary Alexa-conjugated antibodies (S.T. 1) and counterstained with DAPI for 1h at RT, followed by four washes in 1% BSA in PBS. Finally, embryos were mounted and imaged (Nikon Ar1 laser confocal scanning309 microscope (Nikon Eclipse Ti-E) equipped with the NIS-Element 4.40 software).

#### In vitro implantation simulation and Outgrowth assay

To simulate the process of embryo implantation in vitro, an outgrowth test was performed on both TGF-β3-treated and control (CTR) blastocysts. Blastocysts were placed on gelatine-coated plates and cultured in outgrowth medium consisting of DMEM supplement with 10% of fetal bovine serum (FBS), 0.1% glutamine, 0.05% nonessential amino acids, 0.05% gentamicin and 1 μL/mL FGF2.

Each blastocyst (one per well) was trasferred to the the gelatin-coated plates using a mouth capillaries. Since the blastocysts did not attach to the gelatin layer spontaneusly, on day 2 of culture,mechanical implantation was simulated using two fine needles to gently secure the blastocysts onto the surface. The blastocyst were then culture for an additional two days before imaging. The outgrowth area was analyzed by processing the images with ImageJ software to measure the expansion of the blastocyst outgrowth. Embryos were monitored daily, and the medium was refreshed every 2–3 days. Photographic documentation was captured throughout the culture period to track the progression of the outgrowth.

#### Oil Red O staining for lipid Droplet Detection

Lipid droplet accumulation in outgrown cells derived from sheep blastocysts was evaluated using Oil R O staining. Cells cultured in 4-well plate were initially washed with DPBS and then fixed with 4% paraformaldehyde for 30 minutes. After fixation, the cells were stained with 5 μg/mL Oil Red O solution (Sigma, USA) for 10 min, following the protocol decribed by Hou, D. et al. [Hou et al., 2015]. After staining, the cells were carefully rinsed to remove excess stain and were then examined using light microscopy to visualize the presence and distribution of lipid droplets within the cells. Photographs were taken to document the results for further analysis.

### Transcriptome analysis

#### RNA extraction

RNA was isolated from four blastocysts for each replicate and condition by the NucleoSpin miRNA kit (Macherey–Nagel) using the protocol in combination with TRIzol lysis (Invitrogen) and recovery of small and large RNA recovery in one fraction. The quality and quantity of RNA were determined by Agilent 2100. The isolated RNAs were stored at −80°C until use.

#### Library preparation

RNA samples (RIN > 6.5) from three replicates (n =3) for each condition (n=2, control C, Treated with TGF-β3 T) were used for library preparation. RNA-Seq libraries were obtained using the Illumina TruSeq RNA Sample Preparation Version2 kit after polyA enrichment, and libraries enrichement with 22 cycles of PCR, accordingly to what previously reported for sequencing low input translatome samples [Song et al., 2018]. Concentration and quality checks of libraries were determined using Agilent 2100 bioanalyzer. Sequencing was performed on Illumina NovaSeq X, 150 cycles paired end.

#### Data analysis

The nf-core/rnaseq version 3.8.1 pipeline (https://nf-co.re/rnaseq) was run for RNA-Seq data analysis. The pipeline integrates the sequence trimming (TrimGalore version 0.6.7) and sequence alignment (STAR version 2.7.10a), [Dobin et al., 2013]. Sequences were aligned to the Ovis Aries ARS-UI_Ramb_v2.0 reference genome and alignments to gene regions were quantified using Salmon version 1.5.2 (https://combine-lab.github.io/salmon/). Differential expression between samples were calculated with The EdgeR Bioconductor package version 3.6 (Bioconductor, https://bioconductor.org/packages/release/bioc/html/edgeR.html) [Robinson et al., 2010]. Principal Component Analysis PCA and hierarchical clustering were performed with Genesis [Sturn et al., 2002].

#### Quantitative PCR qPCR

4ul of RNA was added of 0.125 ul of random hexamers, 0. 5ul oligodT, 0.5ul dNTPs and 5ul RNAse free water and incubated at 65°C for 5 minutes and placed at 4°C. Mixture was then added of 4 μl RT buffer (5 x), 1μl of DTT, 1ul RNase inhibitor, and 1μl of SuperScript™ II Reverse Transcriptase (Thermo Fisher, Waltham, MA USA). Reverse transcription was carried out at 25°C for 5 minutes, 42°C for 1 hour and 70°C for 15 minutes. The transcript level of the different MX Dynamin Like GTPase 1 (MX1), Inhibin Subunit Alpha (INHA) and Bromodomain and WD Repeat Domain Containing 3 (BRWD3) genes and glyceraldehyde-3-phosphate dehydrogenase (GAPDH), actin beta (ACTB) as a housekeeping gene, were measured using real time-PCR. Supplementary Table reported the primers used in this study. Primer were designed from specific exon-exon junction to avoid DNA genomic. Real-time PCR was performed using 4 μL of diluted cDNA (1:20 Vol.), 5 μl of the SYBR Green Master Mix (Applied Biosystems, Carlsbad, California, USA) and 0.5 μl of forward and reverse primers (final concentration 900nM with QuantStudio 6 Flex Real-Time PCR Systems (Applied Biosystems, Carlsbad, California, USA). Gene relative expression between C and T groups were calculated by the PCR R package [Ahmed & Kim, 2018].

#### Statistical analysis

Quantitative data are expressed as mean±SD. If parametric, Unpaired t test was used to compare the averages/means of two condtions, if not-parametric, Kolmogorov–Smirnov test was used to determine if there is a significant difference between the two groups. Developmental rate were compared by chi-square test. Statistical significance was considered when probability values (p) were lower than 0.05. Statistical analyses were performed using GraphPad Prism software (Version 10.4.1, CA, USA).

## Acknowledgements

We thank all members of involved laboratories for the valuable discussion.

## Author Contributions

Conceptualization: F.B., D.I., P.L., M.C.; Methodology: F.B., D.I., M.C.; Software: E.C., B.L., L.P.; Validation: F.B.; Formal analysis: F.B., L.P.; Investigation: F.B., M.D., M.M.; Resources: M.LS., M.M., M.D., A.S., L.G.; Data curation: D.I.; Writing - original draft: F.B.; Writing - review & editing: R.A., D.I., M.C., P.L.; Visualization: F.B., D.I., R.A.; Supervision: D.I., M.C. ; Project administration: M.C.; Funding acquisition: M.C., D.I., P.L., F.B..

All authors have read the manuscript and agreed with its content

## Data and materials availability

RNA-seq data are deposited in NCBI accession BioProject ID : PRJNA1193800.

Further information and requests for resources and reagents should be directed to and will be fulfilled by the lead contact.

