## Supplementary table1 for "TGF-β3 Promotes Trophoblast Development via ACSS2-Dependent Permissive Lipid Metabolism"

S.T.1 Details of antibodies used immunostaining

| | Target | Host Species | Source | Catalog number | Diluition | |
|---|---|---|---|---|---|---|
| Primary Antobodies | OCT4 | Mouse monoclonal | Santa Cruz | sc-365509 | 1:250 | Used in Fig. 2A |
| | SOX2 | Rat monoclononal | eBioscience | 14-9811-82 | 1:500 | Used in Fig. 5C |
| | CDX2 | Mouse monoclonal | BioGenex | MU392-UCE | 1:50 | Used in Fig. 2A, 5C |
| Secondary Antobodies | Anti-Mouse IgG 488 | Goat | Invitrogen | A11001 | 1:750 | Used in Fig. 2A |
| | Anti-Rat IgG 488 | Goat | Invitrogen | A11006 | 1:750 | Used in Fig. 5C |
| | Anti-Mouse IgG 594 | Goat | Life Technologies Corp. | A21424 | 1:750 | Used in Fig. 2A, 5C |
